# Integrating digital and field surveillance to complement efforts to manage epidemic diseases of livestock: African swine fever as a case study

**DOI:** 10.1101/2021.05.27.445948

**Authors:** Michele Tizzani, Violeta Muñoz-Gómez, Marco De Nardi, Daniela Paolotti, Olga Muñoz, Piera Ceschi, Arvo Viltrop, Ilaria Capua

## Abstract

The SARS-CoV-2 pandemic has unveiled the importance of stakeholders and ordinary citizens in managing infectious disease emergencies. Efficient management of infectious diseases requires a top-down approach which must be complemented with a bottom-up response to be effective. Here we investigate a novel approach to surveillance for transboundary animal diseases using African Swine fever as a model. We were able to collect data at a population level on information-seeking behavior and at a local level through a targeted questionnaire-based survey to relevant stakeholders such as farmers and veterinary authorities. Our study shows how information-seeking behavior and resulting public attention during an epidemic, can be addressed through novel data streams from digital platforms such as Wikipedia. We also bring evidence on how field surveys aimed at local workers (e.g. farmers) and public authorities remain a crucial tool to assess more in-depth preparedness and awareness among front-line actors. We conclude that they should be used in combination to maximize the outcome of surveillance and prevention activities for selected transboundary animal diseases.

## Introduction

African Swine Fever (ASF) is a transboundary animal disease and its impact on global markets can potentially be catastrophic, threatening the economy from local to global level (1).

Despite decades of international control efforts, the disease is still spreading in various regions of the world with different epidemiological dynamics. These are significantly influenced by the natural environment as this influences the density of susceptible species and vectors (i.e. ticks) but also significantly by human behavior (2). In particular, human behavior plays a key role in the transmission and geographic spread of the ASF virus (3,4) through infringement or low compliance with biosecurity measures. Among all, the practice of swill (scraps of meat that are potentially infectious) feeding mainly in backyard farms is known to be a driver of infection. Additionally, underreporting of ASF suspected cases linked to movements of contaminated pork products and infected pigs have contributed to the ASF spread within and across countries (2,4,5). The unavailability of a vaccine or treatment makes awareness programs and mass culling among the few options available for disease prevention and control(5). ASF is a disease of wild and domestic pigs that is present in several European, African, and Asian countries and is heavily threatening their pig industries (1). The World Health Organization for Animal Health (OIE) and the Food Agriculture Organization (FAO) recognize awareness campaigns as essential for prevention and prompt intervention of the health authorities(6).

This complex and heterogeneous scenario calls for novel solutions to preparedness and surveillance that could go beyond traditional public health methodologies. Therefore, in this work, we propose an integrated approach to monitor public awareness and preparedness of animal health authorities and field workers based on a combination of non-traditional data sources and survey-based methodologies. On one hand, assessing public awareness is key to develop effective communication campaigns and prevent the introduction or mitigate the spread of the disease in any given area. Public awareness can be monitored through the exposure of the public to the news which results in proactive information seeking, as measured for example through Wikipedia pageviews. This approach has been already successfully adopted in other contexts, focusing on human diseases (e.g. epilepsy, asthma) (7,8) and some zoonotic diseases, (e.g. Zika, H1N1, and Covid-19)(9–11). On the other hand, knowledge and attitude surveys can be adopted to assess the level of preparedness of local health authorities and field workers who are at the frontline of disease containment and prevention (12–14). Our proposed methodology aims at combining the insight provided by digital data to assess information-seeking behavior and awareness in the general population (with a focus on Europe) with a field survey-based study in Estonia involving pig farmers and local veterinary authorities. We believe this integrated approach, when regularly implemented, could provide better insights on the level of attention of the public towards a specific hazard and risk allowing better communication strategy and ultimately, improving risk management strategies.

ASF was first reported in Estonia in wild boars in 2014 (15) and domestic pigs in 2015 (16). The incursion of the ASF virus to domestic pigs was very likely due to the introduction of contaminated fomites by people (e.g. clothing, vehicles, feed, bedding material) to pig farms as a result of deficient biosecurity measures (16). Estonia has been active in field and research programs on ASF prevention and control and has developed collaboration programs between the farmers and the official veterinary services. The Estonian government has invested efforts in raising awareness of ASF among the general population and animal-related target groups, such as pig farmers and hunters (17,18). Estonia self-declared free of ASF in domestic pig in 2019 (19).

## Material and methods

In this work, we propose a two-fold approach based on digital and stakeholder perspectives by monitoring digital sources, such as news and Wikipedia pageviews as a proxy for information-seeking behavior among the general population as a consequence of news exposure. This activity has the goal of assessing the level of general awareness concerning news coverage of ASF outbreaks. The stakeholder component was assessed through knowledge and attitude field surveys in Estonia focussing on veterinary authorities and farmers to assess the level of preparedness towards ASF.

### Digital news, Wikipedia pageviews, and public awareness

We have collected digital news and Wikipedia click rates for countries in Europe (Belgium, Czech Republic, Estonia, Italy, Latvia, Lithuania, Poland, Romania, Ukraine) and Asia (China, India, South Korea) affected by ASF. Data have been gathered on the period between January 2015 to May 2020. The Wikipedia pageviews information is not available before January 2015. Moreover, we have extracted information about ASF outbreaks from the official reports of the Animal Disease Notification System (ADNS) for European countries (20), and from the OIE update reports on ASF for Asian countries (21).

We used the Wikipedia application programming interface (API) (22) to collect the number of visits per day of Wikipedia articles normalized with the total monthly access to Wikipedia from each targeted country. We selected the Wikipedia articles between 2015/02 and 2020/05 specific for ASF, namely ‘African_swine_fever_virus’, and the relative translations in the languages of interest, (S1 Table). For most of the countries of the study, the language is highly indicative of the location. On the other hand, a weighted normalization factor for the number of views was necessary to account for the multilingualism of some of them, like Belgium and India. More specifically, we weighted the number of daily accesses to a single article from a Wikipedia project (S1 Table) *p, S*_*p*_*(d)*, with the total number of monthly accesses from a country, *c*, to the related Wikipedia project *T*^*c*^_*p*_*(d)*, such that the daily pageviews from a given Wikipedia project and country was described by equation (1):

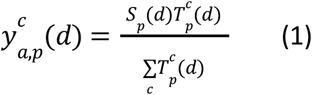

where the denominator is the total number of views of the Wikipedia-specific project. The total volume of views at day, *d*, from a country, *c*, is then given by the sum over all the articles and projects, *p*, given by equation (2):

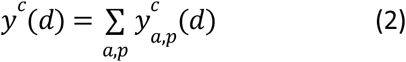

We collected the news from the Global Database of Events, Language, and Tone, (GDELT) project (https://www.gdeltproject.org/), a high-resolution open data platform that monitors news worldwide in over 100 languages, translating and processing them to identify events, people, organizations, locations, themes, languages, and original Uniform Resource Locator (URL) to the article page. We considered all the news published online between 2015/02 and 2020/05 mentioning ASF, for a total of 107,547 articles distributed among the countries as shown in S1 Table, collected using the query in S9. To characterize the signal for each country, we considered the news that was in at least one of the main spoken languages in the country and was mentioning the country either in the location item or in the theme item of the GDELT dataset. Moreover, using the URL we collected and translated to English the summary of the full text of each article to implement a topic modeling analysis.

To analyze the correlation between ASF media coverage in a given country and online users’ collective response as measured through the Wikipedia pageview, we introduced two regression models as shown in Equation number 4. The first one is a simple linear regression, while the second one contains a memory kernel to account for “memory effects” (e.g. loss of interest) in the public response to media coverage. In the latter, the cumulative news articles volume time series were weighted with an exponential decaying term (9,11) introducing the variable in equation (3):

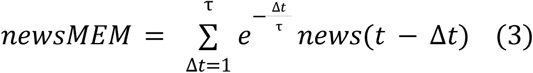

where τ is a free parameter that sets the memory timescale. We tuned τ in the range of [1,60] optimizing the results of the linear regression for the adjusted R^2^, and showing only the best results. Finally, the two models considered are shown in Equation (4):

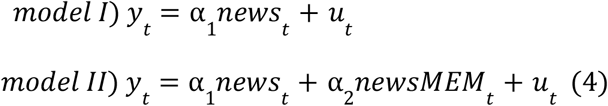

where *y*_*t*_ is the number of country-specific Wikipedia pageviews, and *u*_*t*_ is the error term. More details on the diagnostics for the two models can be found in the Supplementary Information. In the two models, the independent variables are either the news volume or the news volume plus a memory term.

Finally, to qualitatively explore the content of the digital news, we analyze the prevalent topics in the news articles through an unsupervised topic modeling approach (23). Topic modeling is a statistical method that is particularly effective for classifying, clustering, and arranging textual data in latent themes. It has been extensively applied in the literature to extract groups of coherent information from a list of documents(9,23–25). We used a well-known probabilistic framework, the latent Dirichlet allocation (LDA) (26). We cleaned and lemmatized the text using the python “spacy” library (27) while the number of topics was chosen through a grid search of the parameter for the Latent Dirichlet Algorithm from the scikit-learn python library (28). The process led originally to twenty-five topics which were lately grouped into 5 main broad topics. All the authors (the majority of which have a background in veterinary sciences) were involved in the qualitative annotation of the resulting topics. The news analyzed in this activity was focused on Estonia to qualitatively assess the coverage of the problem in this specific country where we have also carried out more in-depth analysis by means of field surveys described in the next section.

### Surveys in Estonia

The main aims of the surveys in Estonia were to explore the use of social media by pig farmers, to collect information on the ASF communication strategy of veterinary authorities, and to explore the perception of the veterinary authorities on the level of ASF awareness in Estonian farmers and hunters. Estonia was selected as a case study due to its advanced digital environment (29) and due to the recent circulation of ASF in both wild boar and domestic pig populations(16). Two questionnaires were designed, one targeting pig farmers and one targeting risk managers within the veterinary authorities. Both questionnaires were designed and piloted in English and can be found in S3 and S4 of the supporting material. The questionnaire for pig farmers was translated into Estonian by collaborators from the Estonian University of Life Sciences (Estonia) and aimed at collecting information on demographics, type of production, experience, and perceptions related to ASF, use of social media, and perception of the implementation of the ASF risk management strategy, using closed-ended questions. The questionnaire for pig farmers was delivered at the end of five focus group discussions, organized and implemented by the Estonian University of Life Sciences (Estonia), where farmers were asked to complete the questionnaire independently. The focus groups were divided into two sessions: the first session aimed at assessing the level of existing knowledge about clinical signs, transmission routes, and preventive control measures while the second session aimed at exploring participants’ attitudes towards the consequences during and after an ASF outbreak in domestic pigs.

The questionnaire for the veterinary authorities, formed by open-ended questions was delivered in English, and it was designed to collect information on the coordination practices related to the national risk management of ASF, the involvement of farmers and wild boar hunters in risk management strategies, and the national communication strategy on ASF. The questionnaire was delivered through both face-to-face and phone structured interviews. Since the survey did not include active sampling in animals/humans and no personal information of respondents was collected, it was not necessary to request Ethics committee approval. Before starting the interview, a confidentiality statement was agreed upon with all participants and after given consent, the interview was conducted. The data collection process for both target groups took place from November 2019 to February 2020 and it was interrupted due to COVID-19 restriction.

Survey data were gathered in Excel ®. A descriptive analysis of all the questions was conducted for both questionnaires. Outputs related to few questions (not relevant to the topics covered in the current study) will be described in a separate manuscript under preparation. Questions that are included in this study are indicated in the questionnaires in the supplementary documentation.

## Results

### Digital news, Wikipedia pageviews, and public awareness

In Figure 1 we show the comparison between weekly aggregated normalized volumes of news and Wikipedia pageviews, for each country compared with the reports of ASF outbreaks in the selected countries for domestic pigs and wild boars.

**Figure 1.**
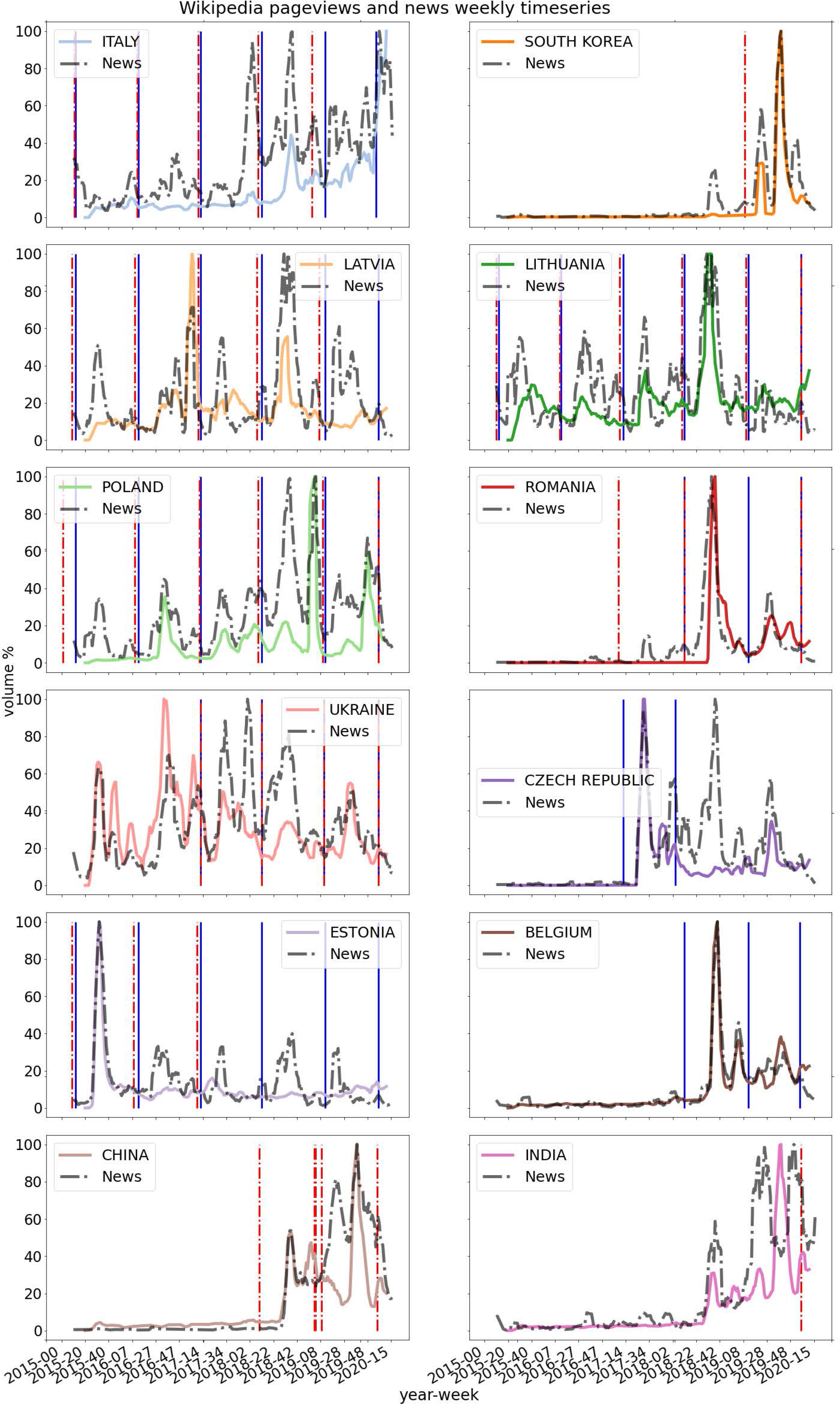
Weekly aggregated data of Wikipedia page view count, colored-solid lines, News volume, dotted-black lines, and ASF surveillance cases reports for domestic pigs, dashed-red vertical lines, and wild boars, solid blue vertical lines.

The temporal profile of the two digital signals was very similar, showing synchronization of the time-series for most of the countries, as confirmed by correlations in Table 1. Notice that the information-seeking behavior decreased after having reached the highest peak, even if the exposure to the news remained high. This was particularly evident in Estonia, Romania, Belgium, and the Czech Republic. Also, Latvia, Lithuania, and Poland had similar temporal profiles, presenting multiple peaks in both Wikipedia pageviews and the volume of news during the reporting time of the ASF outbreaks. The communication of official reports of the outbreaks was not always reflected in the news volume, failing to trigger an information-seeking behavior towards the subject. This was particularly evident in Italy where, despite multiple ASF cases being reported before October 2018, Wikipedia searches only started after the first peak of news volume. These trends indicated a necessary threshold of information exposure to trigger the information-seeking behavior of the general population, which was not necessarily dependent on the surveillance reports of the outbreaks.

**Table 1.** Pearson and Spearman correlation between Wikipedia pageviews and news volume.

| Wikipedia – News by source country | Pearson | Spearman |
| --- | --- | --- |
| ITALY | 0,55 | 0,6 |
| SOUTH KOREA | 0,75 | 0,7 |
| LATVIA | 0,42 | 0,22 |
| LITHUANIA | 0,53 | 0,23 |
| POLAND | 0,53 | 0,72 |
| ROMANIA | 0,64 | 0,71 |
| UKRAINE | 0,33 | 0,46 |
| CZECH REPUBLIC | 0,57 | 0,79 |
| ESTONIA | 0,65 | 0,23 |
| BELGIUM | 0,86 | 0,71 |
| CHINA | 0,74 | 0,73 |
| INDIA | 0,4 | 0,72 |

In Table 2 we show the results of the two regressions models and in Table 3 we show the coefficients estimates for the model with the memory parameter.

**Table 2.** Adjusted R^2^ for the two linear regression models applied to predict Wikipedia visits.

| Countries | Model I | Model II |
| --- | --- | --- |
| ITALY | 0.57 | 0.72 |
| SOUTH KOREA | 0.57 | 0.73 |
| LATVIA | 0.44 | 0.59 |
| LITHUANIA | 0.46 | 0.53 |
| POLAND | 0.36 | 0.37 |
| ROMANIA | 0.47 | 0.59 |
| UKRAINE | 0.51 | 0.61 |
| CZECH REPUBLIC | 0.43 | 0.48 |
| ESTONIA | 0.56 | 0.61 |
| BELGIUM | 0.78 | 0.79 |
| CHINA | 0.65 | 0.65 |
| INDIA | 0.28 | 0.64 |

**Table 3.** Coefficients estimates for the model with memory. The coefficients are all significant with *p<0*.*001*, except for Polonia (*p=0*.*009*), Romain (*p=0*.*013*), Belgium (*p=0*.*003*), and China (*p=0*.*02*).

| Countries | News | News+Memory |
| --- | --- | --- |
| ITALY | 0.20 [ 0.03, 0.37] | 0.14 [ 0.11, 0.17] |
| SOUTH KOREA | 0.41 [ 0.16, 0.65] | 0.43 [ 0.19, 0.68] |
| LATVIA | 0.17 [ 0.00, 0.34] | 0.08 [ 0.06, 0.11] |
| LITHUANIA | 0.41 [ 0.04, 0.78] | 0.09 [ 0.04, 0.13] |
| POLAND | 0.29 [-0.05, 0.62] | -0.05 [-0.21, 0.11] |
| ROMANIA | 0.19 [-0.13, 0.51] | 0.25 [ 0.16, 0.35] |
| UKRAINE | 0.30 [ 0.14, 0.46] | 0.34 [ 0.25, 0.43] |
| CZECH REPUBLIC | 0.25 [ 0.07, 0.43] | 0.16 [ 0.04, 0.27] |
| ESTONIA | 0.25 [ 0.03, 0.48] | 0.16 [ 0.10, 0.23] |
| BELGIUM | 0.61 [ 0.36, 0.85] | 0.09 [-0.03, 0.20] |
| CHINA | 0.49 [ 0.17, 0.80] | -0.07 [-0.22, 0.09] |
| INDIA | -0.07 [-0.23, 0.08] | 0.36 [ 0.20, 0.53] |

We compared the two models by using the adjusted coefficient of determination (R^2^) (Miles, 2005). We found that memory effects improve the model performance. Additionally, we compared the two models using the F-test for nested models (30), obtaining *p<0*.*001* in most of the cases except for Polonia, Belgium, and China, for which *p<0*.*02*. Hence, strong statistical evidence suggested that adding the memory term improves the performance.

Finally, Figure 2 shows the main topics in the Estonian news dataset. The most relevant one/ones refer/refers to control measures, intended to prevent both the spreading of ASF and the presence of infected products on the market. This result confirms the most prevalent exposure of the Estonian population to information about control measures during the entire period of the study. Although we cannot classify which proportion of the reached population are farmers or authorities, we can assume, given the specificity of the ASF topic, that they are a part of it. In this case, the exposure to control measures information was confirmed also by the survey of the veterinary authorities.

**Figure 2.**
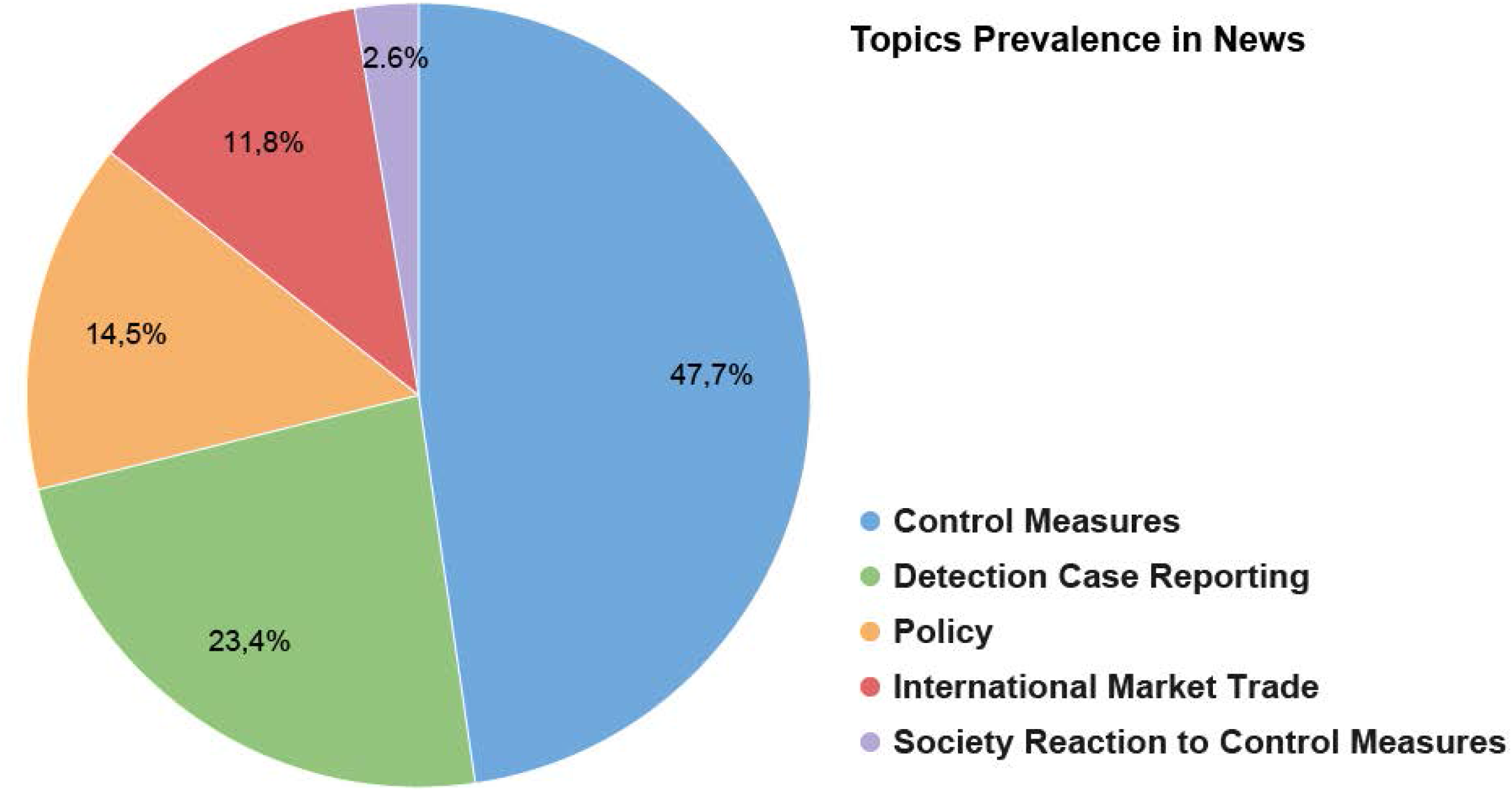
Percentage of documents per topic.

### Questionnaires in Estonia to pig farmers and veterinary authorities

A total of 22 pig farmers answered the questionnaire. Demographic information regarding the farm’s counties can be found in (S5 Table). Out of the 22 respondents, 4(18.2%) had a multiplier production type, 7(31.8%) kept fattening pigs and 11(50%) manage a farrow-to-finish farm type. A total of 18 (81.8%) farmers declared that ASF has never been detected in their farms against 4(18.2%) that mentioned that it was detected. Also, 17(77.3%) farmers felt confident in recognizing the clinical signs of ASF in pigs and 5(22.7%) did not. All farmers mentioned that they have invested resources in improving the biosecurity of their farms as a result of the ASF epidemic in Estonia. The five main biosecurity aspects in which farmers have invested resources included training staff on disease management (20/22, 90.9%), fencing (20/22, 90.9%), cleaning and disinfection (18/22, 81.8%), management of dead animals (17/22, 77.3%) and transport of animals (17/22, 77.3%). S6 Table in the supplementary material contains all the biosecurity measures in which Estonian farmers invested resources due to ASF. All farmers were aware that ASF is currently spreading in other countries. The five most selected sources to access information on ASF were social media (14/22, 63.6%), other internet sources (non-social media) (13/22, 59.1%), colleagues/friends (13/22, 59.1%), agricultural magazines (12/22, 54.5%), general papers/magazines (12/22, 54.5%). Farmers were also asked about the use of social media as a tool to share information on topics related to ASF. From a predefined list of possible answers provided, half of the farmers (11/22, 50%) reported that social media is useful to share information on ASF related topics, eight farmers (36.4%) mentioned that they prefer using other communication channels, and three (13.6%) that they do not normally use social media.

Regarding the farmers’ perception of the implementation of the ASF strategy, the majority of farmers (21/22, 95.5%) felt well-informed about the measures recommended by the veterinary authorities to prevent/control ASF outbreaks in pigs. Similarly, most farmers (17/22, 77.3%) thought that there is cooperation between the farmers’ community and the veterinary authorities. Also, 15 out of the 22 farmers (68.2%) thought that farmers are sufficiently consulted by the veterinary authorities in terms of ASF control and 17(77.3%) agreed with the majority of the measures to prevent/control outbreaks of ASF in pigs. Out of the 22 farmers, 17 (77.3%) agreed with the majority of the measures to prevent/control outbreaks of ASF in pigs and 5 (22.7%) did not. However, 16 out of 22 farmers (72.7%) did not feel satisfied with the economic support from the government to control ASF.

We assessed the level of satisfaction of the farmers (very dissatisfied, slightly dissatisfied, slightly satisfied, and very satisfied) with the veterinary and other veterinary authorities concerning ASF communication practices considering two aspects, the communication promptly to farmers on ASF-related topics during an outbreak and the content of the communication to farmers on ASF-related topics during an outbreak. About fifty-four percent (54.5%, 12/22) of respondents were slightly satisfied, 27.3% (6/22) slightly dissatisfied, 13.6% (3/22) very satisfied, and 1 respondent very dissatisfied with the timely manner of the communication to them on ASF-related topics during an outbreak. Half of the respondents (11/22, 50%) were slightly satisfied and 31.8% (7/22) were slightly dissatisfied and 18.2% (4/22) were very satisfied with the content of the communication on ASF-related topics directed to farmers during an outbreak. Also, 45.5% (10/22) of respondents felt slightly dissatisfied, 36.4% (8/22) slightly satisfied, 9.1% (2/22) very dissatisfied, and one respondent very satisfied with the farmers’ involvement of the farmers’ community in the national ASF control strategy. Regarding the implementation of actions by the veterinary and other veterinary authorities promptly during an ASF outbreak, 45.5% (10/22) felt slightly satisfied, 31.8% (7/22) slightly dissatisfied, 13.6% (3/22) very satisfied, and 9.1% (2/22) very satisfied.

Eleven interviews were conducted with representatives of the Estonian veterinary authorities, three working at the central level and eight respondents working at the district level for the Estonian competent authorities. Interviewees were asked which main institutions and private organizations were involved in the risk management strategy of ASF in domestic pigs and wild boars in Estonia. S7 Table, in supplementary material, shows the name and the number of nominations of public institutions and private organizations mentioned by respondents. Interviewees were asked about which target groups, communication channels, and materials are used as part of the ASF communication strategy. Their answers can be found in S8 Table in the supplementary material. The coordination between public institutions and private organizations was described as good for most respondents (8/11, 72.7%), adding that it has improved from the beginning of the epidemic until the moment that the interview was taken. Most of the barriers encountered in the coordination mentioned by respondents were related to communication aspects (47.4%), lack of direct funding to support coordination (15.8%), poor commitment (15.8%), lack of clear chain of command (10.5%) and lack of transparency (10.5%).

The coordination between the district and central veterinary authorities was positively assessed by respondents (9/11, 81.8%). Only one respondent assessed this coordination as dissatisfactory and other respondents did not answer this question.

## Discussion

In experiencing the COVID 19 pandemic, it is clear that epidemic diseases are to be managed at the general population level as well as at the local level, and that these two approaches must be synergic and have a common goal. ASF is a perfect example to investigate as disease control measures rely on combating the risk of introduction and spread by applying measures at a farm level, at the national level, and the level of the general population. Citizens are being recommended on proper individual behaviors (e.g. proper disposal of food in recreational areas) and help implementation of passive surveillance (e.g. reporting of wild-boar carcasses from hunters and tourists to authorities). In this study, we assessed the information-seeking behavior as well as the awareness and the preparedness of ASF at two different scales (general population and target groups) intending to evaluate the applicability of an integrated approach to monitor public awareness and preparedness of health authorities and field workers based on a combination of non-traditional data sources and survey-based methodologies.

In the case of ASF, specific sub-groups of the population that are at the forefront of an epidemic preparedness, such as pig farmers, differ from the general population in the sense that they are considerably more aware and interested in a topic that affects them directly, and this is reflected in the results of the questionnaire. Not the same can be said for the general population, given that digital information-seeking behavior peaked only after generalist news coverage. For these reasons, both approaches (digital and field-oriented) are valuable and needed from a public health perspective to understand how interest, risk perception, and awareness in the general population and specific interest groups evolve during an epidemic and how they may affect public opinion as well as the preparedness.

First, we exploited the pervasiveness of digital data to assess the awareness among the general population as measured through Wikipedia page views in reaction to the exposure to news about ASF outbreaks at an international scale. Then, we examined the preparedness, awareness, and information-seeking in localized areas through questionnaires to farmers and health authorities, with a focus on the Estonian context.

Expectedly our results show that, at an international scale, public interest rapidly declines after an initial attention peak which occurs after exposure to news coverage of a specific outbreak. In particular, the public activity profile as measured through the access to the Wikipedia pages shows nonlinear dependencies and memory effects in the relation between information seeking, media pressure, and disease dynamics.

This suggests that governmental and health authorities outreach strategies should be dynamic and customized to the evolving level of attention of the public. This result has important public health implications. Consistently with previous literature work for different human diseases and zoonotic outbreaks(9,11), we found that Wikipedia page activity is highest in correspondence of media coverage of official communications by public health authorities. This is relevant because official communications amplified by media coverage could effectively be leveraged to capture the public attention and stimulate information-seeking even on a longer-term after the outbreak. This is particularly true in the post-Covid-19 environment which has primed the general public to the importance of individual behavior in the spread of epidemic diseases of humans and animals(9).

Moreover, as shown in the topic discovery activity, news articles covered a broad range of topics: control measures, detection, case reporting, and policy. In particular, these are the main topics on the news coverage in Estonia in addition to “society reaction to control measures”. This points to the fact that, in the specific case of Estonia, the media have given visibility to population groups specifically affected by the problem of ASF.

The focus on Estonia has been further explored through the field data collection (surveys proposed to farmers and veterinary authorities) which allowed us to assess communication practices in a country that was heavily affected by the disease (31). The majority of farmers felt well-informed about the measures recommended by the veterinary authorities to prevent/control ASF outbreaks in pigs. Similarly, most farmers thought that there is cooperation between the farmers’ community and the veterinary authorities. This is a finding of relevance because usually farmers tend to complain against veterinary authorities because of culling practices and poor subsequent compensations (32,33). In our sample, despite having 16 out of 22 farmers not satisfied with the compensation, the majority was still positive about the risk management strategies implemented. Moreover, farmers reported being satisfied with the timeliness of communication on ASF by the health authority during an outbreak. This points to a positively evolving relationship as, in general, there is the perception among farmers that the authority could have been more efficient when it comes down to infectious disease management (34).

Importantly, the coordination between public institutions and private organizations was described as good for most respondents, adding that it has improved from the beginning of the epidemic until the moment that the interview was taken. The coordination between all these actors (public and private) is known to be crucial for disease control. Hunters, farmers, private veterinaries associations are key players in ASF risk management strategies and, therefore, also coordination among these actors is extremely relevant. Our study shows that also timeliness of transmitting official communication between districts and the central level during an ASF outbreak was referred to as “satisfactory” by about half of the respondents. This is relevant since the epidemiological situation is evolving and may be subject to change and thus require revised control strategies. Finally, almost all the respondents mentioned that there is cooperation between farmers/hunters and the veterinary authorities to control ASF and this certainly has facilitated keeping ASF mostly under control in Estonia.

On the other hand, among the compliance issues mentioned by the health authority respondents (lack of financial support and knowledge, cooperation and commitment of farmers and hunters, lack of detailed ASF legislation, lack of regular training and equipment), lack of support from the press was also mentioned. This is in line with the information-seeking behavior that we observed by looking at the amount of news devoted to outbreaks and the expected declining attention level from the general public once the news coverage is over.

Finally, the local health authorities identified difficulties in addressing specific preventative aspects which are related to ASF such as reaching moving target groups (e.g. travelers, truck drivers) and keeping them engaged. These are important categories in the general population to reach out to as in previous experience travelers/truck drivers, foreign workers were suspected of being involuntary involved in the introduction of the disease in Belgium (35). This is also in line with the lack of public attention detected with digital data.

### Limitations

During the study, we faced several limitations, especially with field activities which were complicated by the concurrent Covid-19 pandemic on multiple fronts. These included the small sample size of both Estonian farmers and veterinary authorities. Only 22 out of 30 farmers (the initial target) have responded to the questionnaire. The disruption of the data collection process was caused to COVID-19 restrictions on travel hampering the implementation of the survey.

A way to overcome the logistic limitations imposed by field surveys, the implementation of digital surveys for both sectorial workers and the general population could bring considerable benefits. This would extend the reachable countries and provide guidelines for the implementations of optimal country-specific strategies to improve awareness for both sectorial workers and the general population.

Also, the fact that Estonian pig farmers filled in the questionnaire after focus group discussions on ASF control measures might have brought anchoring bias, affecting their answers to the questionnaire.

### Future work

Possible future directions for this work would focus on creating a friendly – user digital version of the surveys (i.e. using R-Shiny (36) or similar) that could be regularly used by authorities aimed at reaching out to a larger number of farmers and health authorities in a multi-country fashion. Along these lines, and concerning the evolving epidemiological situation -developing a format that could allow “living questionnaires” would represent an innovative tool to manage established epidemic situations. This could be relevant for ASF given that ASF notifications do not appear to be declining in many countries but could also apply to many other hazards with relevant public health impacts. This could also represent a means of engaging the population categories which can be affected by the problem of ASF both through content delivery and to encourage active participation in the collection of information.

## Acknowledgments

IC acknowledges the Leonardo Fellowship Program, and the Circular Health Initiative at the University of Florida for supporting the work. DP and MT acknowledge the support from the Lagrange Project of the Institute for Scientific Interchange Foundation (ISI Foundation) funded by Fondazione CRT of Torino. MDN and VM acknowledge the support of the Zurich Insurance Company for funding the fieldwork in Estonia.

All the authors thank R Gollakner, SE Stefanou, T Denagamame, Wojciech Iwaniak, Lyudmyla Marushchak, and C Manes for their help and support in gathering and translating information from selected countries.

## Supporting information

**S1 Figure.**
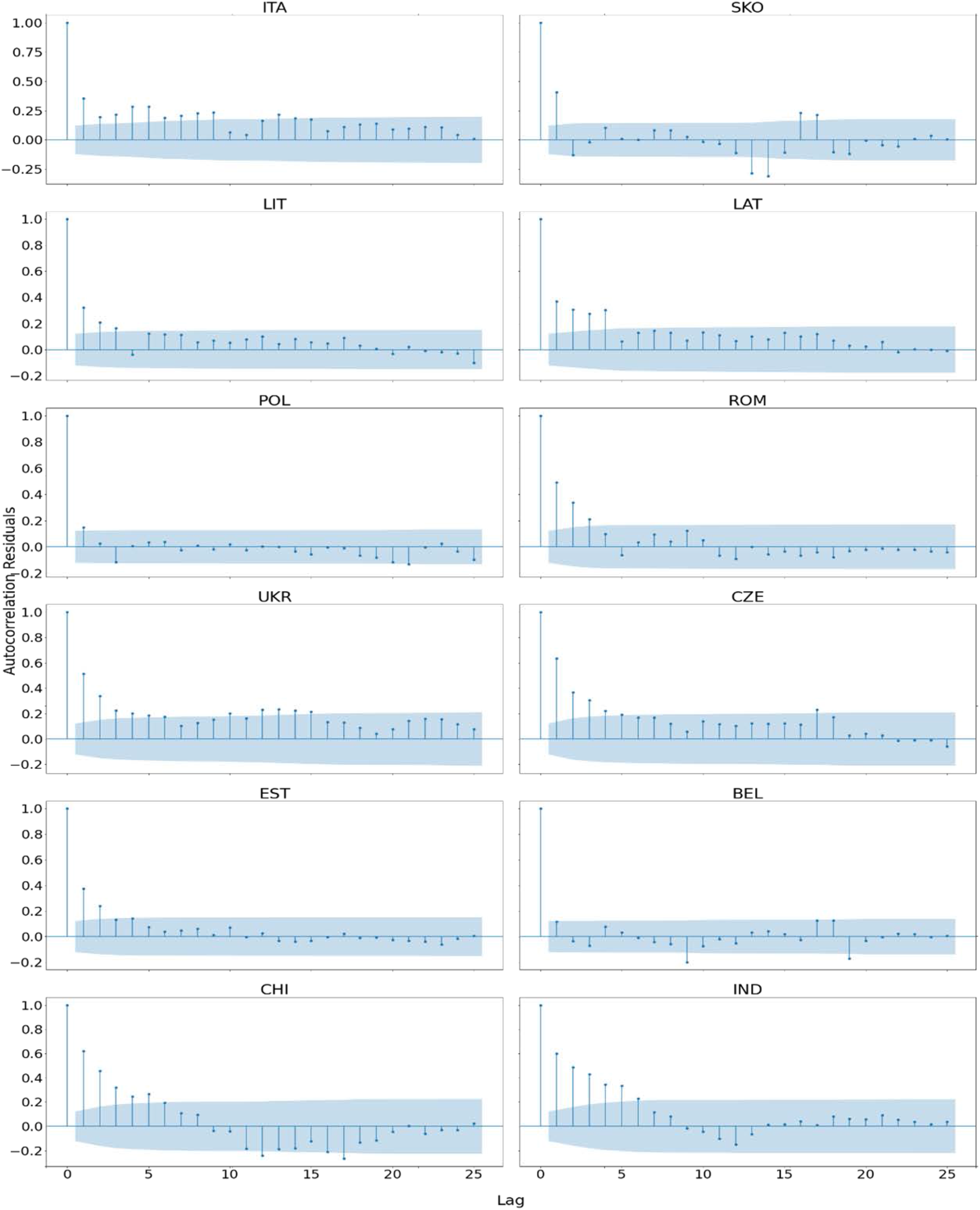
Autocorrelations by country, Model II. Linear Regression-Diagnostics

S1 Table. Total pageviews and news by selected country.

S2 Table. P-values for the Breusch-Pagan Lagrange Multiplier test on heteroscedasticity. The null hypothesis is that residuals are homoscedastic, hence a p-value < 0.05 indicates heteroscedasticity

S3 Questionnaire to Estonian farmers. Selected questions analyzed in this study are marked with a star*

S4 Questionnaire to the Estonian veterinary authorities. Selected questions analyzed in this study are marked with a star*

S5 Table. Number of farmers interviewed by county (Estonian region).

S6 Table. Biosecurity aspects that Estonian farmers mentioned having invested resources due to ASF.

S7 Table. Public institutions and private organizations mentioned by the Estonian veterinary authorities as part of the risk management strategy.

S8 Table. Target groups, information material, and communication channels mentioned by the Estonian veterinary authorities.

S9 GDELT query for data collection.

