## Supporting materials for "Integrating digital and field surveillance to complement efforts to manage epidemic diseases of livestock: African swine fever as a case study"

### Supporting material

S1 Table. Total pageviews and News by selected country.

| Country | Total Pageviews | Wikipedia Project | Total News |
|---|---|---|---|
| Italy | 37758 | it.wikipedia.org | 1143 |
| Lithuania | 9776 | lt.wikipedia.org | 7398 |
| Latvia | 10318 | lv.wikipedia.org | 1167 |
| Romania | 84998 | ro.wikipedia.org | 3974 |
| Ukraine | 126303 | uk.wikipedia.org | 3006 |
| Czech Republic | 25796 | cs.wikipedia.org | 2491 |
| Estonia | 18053 | et.wikipedia.org | 916 |
| Belgium | 6839 | fr.wikipedia.org, de.wikipedia.org | 1300 |
| Poland | 321651 | pl.wikipedia.org | 2861 |
| India | 4856 | en.wikipedia.org, hi.wikipedia.org | 35128 |
| China | 7864 | zh.wikipedia.org | 2423 |
| South Korea | 11514 | ko.wikipedia.org | 1143 |

S2. Linear Regression – Diagnostics

In the main text, we use a Liner Model to nowcast the amount of Wikipedia page views media coverage as independent variables. Here, we test model assumptions of homoscedasticity and no-autocorrelation of residuals. Homoscedasticity of residuals is assessed with the Breusch-Pagan Lagrange Multiplier test (Breusch & Pagan, 1979) on a fitted Ordinary Least Square model (OLS) using python statsmodels library (https://www.statsmodels.org/). Assuming a significance level $\alpha = 0.05$, we obtain heteroscedasticity of residuals for Model II and Model III (S2 Table).

S2 Table. P-values for the Breusch-Pagan Lagrange Multiplier test on heteroscedasticity. The null hypothesis is that residuals are homoscedastic, hence a p-value < 0.05 indicates heteroscedasticity.

| Countries | Model I | Model II |
|---|---|---|
| Italy | 0.0 | 5.3E-07 |
| South Korea | 0.0 | 6.7E-35 |
| Lithuania | 0.0 | 1.2E-09 |
| Latvia | 0.0 | 1.9E-21 |
| Poland | 0.0 | 3.9E-17 |
| Romania | 0.0 | 1.1E-21 |
| Ukraine | 0.0 | 4.4E-09 |
| Czech Republic | 0.0 | 9.8E-11 |
| Estonia | 0.0 | 2.7E-13 |
| Belgium | 0.0 | 1.0E-17 |
| China | 0.0 | 4.1E-21 |
| India | 0.0 | 1.2E-10 |

We check for the autocorrelation of residuals in S1 Figure, where we plot the partial auto-correlation function for Model II (analogous results are obtained for Model I). We notice a significant correlation at the first four lags in all cases, and in some cases also at higher orders.

S3 Questionnaire to Estonian farmers. Selected questions analysed in this study are marked with a star*

**SECTION 1: Demographic information**

1. *Give the name of the County and City/Village where your farm is located.
   County_________
   City/Village _________

**SECTION 2: Farm characteristics**

2. *What is your type of production? Select one answer.

   - o (1)Multiplier[1]
   - o (2)Fattening[3]
   - o (3)Farrow-to-finish[2]

3. How many pigs of each type do you keep in your farm? Select one answer for each type of pigs.

| | | | | |
|---|---|---|---|---|
| Sows | o 1-10 | o 11-100 | o 101-1000 | o >1000 |
| Weaned pigs | o 1-10 | o 11-100 | o 101-1000 | o >1000 |
| Fattening | o 1-10 | o 11-100 | o 101-1000 | o >1000 |

**SECTION 3: Experience related to African Swine Fever (ASF)**

4. *Has ASF ever been detected in your farm? Select one answer.

   - o No
   - o Yes
   - o I do not know

5. *Do you feel confident that you can recognise the clinical signs of ASF in pigs? Select one answer.

   - o No
   - o Yes

6. *Have you invested resources in improving the biosecurity of your farm (equipment, practices, vehicles, facilities etc.) as a result of outbreaks of ASF? Select one answer.

   - o No (Go to question 8)
   - o Yes (Go to question 7)

7. *In what aspects of biosecurity did you invest resources in your farm due to ASF?

[1] Multiplier: Pig herd formed by breeding animals and piglets (up to weaned pigs)
[2] Farrow-to-finish: Pig herd that comprises all the categories of pigs.
[3] Fattening: Pig herd consisted of fattening pigs.

□ (1)Feed, water
□ (2)Equipment supply
□ (3)Transport of animals
□ (4)Removal of manure
□ (5)Fencing
□ (6)Sourcing of animals
□ (7)Management of dead animals
□ (8)Training staff on disease management
□ (9)Cleaning and disinfection
□ (10)Constructional changes
□ (11)Other____________________

8. Which of the following statements applies to you since ASF started spreading? Please, select all that apply.

□ (1)Due to the spread of ASF, I changed the production type of my farm (e.g. from multiplier to fattening pigs)
□ (2) Due to the spread of ASF, my working routine has been affected.
□ (3) Due to the spread of ASF, I am more alert on the health status of my pigs.
□ (4) I do not care about ASF.
□ (5)Other:__________

9. Which of the following statements reflects your perception of the price of pork meat since ASF started spreading? Select one answer.
   o (1)The price of pork meat at farm gate has increased.
   o (2)The price of pork meat at farm gate has decreased.
   o (3)The price of pork meat at farm gate has remained unchanged.

10. *Do you think that ASF is currently spreading in other countries? Select one option.

   o No (Go to question 12)
   o Yes (Go to question 11)

11. *Where did you learn this? Check all that apply.

□ (1)Social media[4]
□ (2)Other internet resources (non-social media)
□ (3)Colleagues/friends
□ (4)Local news media (TV –radio)
□ (5)General papers/magazines
□ (6)Agricultural magazine
□ (7)Other______________

12. *What do you think about the use of social media to share information on topics related to ASF? Select all that apply.
   □ (1)Social media are useful to share information on topics related to ASF.
   □ (2)I prefer sharing information on ASF using other communication channels.

[4] Social media: websites and applications to share information and be part of social networking (e.g.Facebook, Instagram, LinkedIn etc...)

- ◻ (3)I do not normally use social media so I do not have an opinion about this question.

13. Do you think that pig farms in the country are threatened by ASF at the moment?
    - ◻ Yes
    - ◻ No

**SECTION 4: Perception of the implementation of the strategy on African Swine Fever (ASF)**

14. Select one answer for each of the following questions:

| | | |
|---|---|---|
| ▪ *(1)Do think that there is cooperation between the farmers' community and the authorities to prevent/control ASF? | o Yes | o No |
| If No, please, specify briefly why: | | |
| ▪ *(2)Do you think that farmers are sufficiently consulted by the authorities when it comes to prevent/control ASF? | o Yes | o No |
| If No, please, specify briefly why: | | |
| ▪ *(3)Are you satisfied with the economic support from the government to control ASF? | o Yes | o No |
| If No, please, specify briefly why: | | |
| ▪ *(4)Do you feel you are well-informed about the measures recommended by the authorities to prevent/control outbreaks on ASF in pigs? | o Yes | o No |
| If No, please, specify briefly why: | | |
| ▪ *(5)Would you say that you agree with the majority of the measures to prevent/control outbreaks on ASF in pigs? | o Yes | o No |
| If No, please, specify briefly why: | | |

15. What is your level of satisfaction with the veterinary and other authorities in relation to ASF? Tick one option for each of the following aspects.

| Aspects: | Very satisfied | Slightly Satisfied | Slightly Dissatisfied | Very dissatisfied |
|---|---|---|---|---|

| | | | | |
|---|---|---|---|---|
| *(1)Involvement of the farmers' community in the national ASF control strategy | o | o | o | o |
| *(2)Communication in a timely manner to farmers on ASF-related topics during an outbreak | o | o | o | o |
| *(3)Content of the communication to farmers on ASF-related topics during an outbreak | o | o | o | o |
| *(4)Implementing actions in a timely manner during an ASF outbreak | o | o | o | o |

S4 Questionnaire to the Estonian veterinary authorities. Selected questions analysed in this study are marked with a star *

**SECTION 1: General**

1. *First, could you please tell me about your **job position** and **responsibilities** in your organization when it comes to ASF?
   *Please, tell me more…*

**SECTION 2: National risk management strategy of African Swine Fever (ASF)**

2. Could you please tell me **what are the main risk management strategies** targeting domestic pigs (DP) and wild boars (WB) in Estonia?

3. Do you know if these strategies are **following a specific EU regulation**? Could you please tell me which one?

4. Regarding the measures that you mentioned earlier as part of the risk management strategy, do you know if there is a/some measures that is/are implemented in Estonia but that are not part of the EU recommendations?

5. *Given the epidemiological situation of ASF in Estonia, would you say that the incidence of ASF in DP and WB is….*

   a. *Decreasing*
   b. *Stable*
   c. *Increasing*
   d. *Epidemic* *Why do say that…?*

**SECTION 3: Coordination practices related to the national risk management of African Swine Fever (ASF)**

6. *Could you please tell me which are the **main institutions and organizations (public and private)** actively involved in the risk management strategy of ASF (in domestic pigs (DP) and wild boars (WB) in Estonia?

7. *How would you **describe the coordination** between these institutions? From your perspective, which are the **main barriers** encountered in the coordination?
   *Why do you say that…?*

   *Potential barriers to mention:*
   a. *Communication*

b. *Transparency*
c. *Lack of clear chain of command*
d. *Poor commitment from people at different level*
e. *Language (use of terminology)*
f. *Lack of direct funding to support coordination*
g. *Others. Please add:…………………………………………*

8. *How would you assess that is the **coordination** between the regional/**district** and national/**central veterinary authorities**?
   a. *Very satisfactory*
   b. *Slightly satisfactory*
   c. *Satisfactory*
   d. ***Slightly dissatisfactory***
   e. ***Dissatisfactory***
   f. ***Very dissatisfactory***

   *In case of A/B/C,*
   - *What specifically do you find satisfactory?*

   *In case of D/E/F,*
   - *What specifically do you find dissatisfactory?*

9. How would you <u>describe</u> the **timeliness of response** of a/b/c…….between the district and central level during an ASF outbreak (in WB/DP)?
   a. *Transmitting official communication*
   b. *Transmitting informal internal communication*
   c. *Implementing control measures*

<u>Timeliness</u>: *well-timed; the fact of happening at the best possible time or at the right time*

*Why do you say that?*

From your perspective, would you say that the timeliness response is for a/b/c …

d. *Very satisfactory*
e. *Satisfactory*
f. *Dissatisfactory*
g. *Very dissatisfactory*

*Why do you say that…?*

**SECTION 4: Involvement of farmers and wild boar hunters in risk management strategies**

Since ASF started spreading in Estonia in 2014 in WB and in 2015 in DP (Nurmoja *et al.*, 2018), the measures taken by the Estonian authorities to face ASF in the country have been changing throughout the time.

10. What would you say about the **level of involvement** of farmers/ wild boar hunters in those risk management practices to face ASF?

11. Do you think that **there is cooperation** between farmers/hunters and the veterinary authorities to control ASF?

    *Why do you say that…please, tell me more.*

    ***If No**, please, specify briefly why*

12. Do you think that farmers/hunters are **sufficiently consulted** by the authorities when it comes to prevent/control ASF?

    *Why do you say that….please, tell me more.*

    ***If No**, please, specify briefly why*

13. Would you say that farmers are **economically well-supported** when it comes to control ASF?
    *Why do you say that…please, tell me more.*

14. Do you feel that farmers/hunters are **well-informed about** the measures recommended by the authorities to prevent/control outbreaks on ASF in pigs?

    *Please, expand you answer a bit more.*

15. How do you rate the **overall performance** (mainly focus on two aspects formal compliance, purpose of the rules achieved) of farmers/hunters in terms of:

*Formal compliance- with this I mean if they act obeying the orders or rules.*

*Purpose achieved-with this I mean if the intention behind of the rules are achieved or not…)]*

*Four levels for rating: poor, fair, good, excellent…*

| **FARMERS** | **Formal compliance** | | | | **Purpose achieved** | | | |
|---|---|---|---|---|---|---|---|---|
| | Poor | Fair | Good | Excellent | Poor | Fair | Good | Excellent |
| Reporting suspicious cases of ASF in pigs (including death animals) | | | | | | | | |
| Implementation of adequate biosecurity measures | | | | | | | | |

| **HUNTERS** | **Formal compliance** | | | | **Purpose achieved** | | | |
|---|---|---|---|---|---|---|---|---|
| | Poor | Fair | Good | Excellent | Poor | Fair | Good | Excellent |
| Reporting WB found dead | | | | | | | | |

| |
|---|
| Reporting suspicious cases of ASF in WB |
| Implementation of adequate biosecurity measures during transport and slaughter of WBs |

16. Do you think that this compliance/purpose achieved have **changed** throughout the time? From your perspective, which have been the **main barriers** encountered?
    *Why do you say that…?*

**SECTION 5: National communication strategy on African Swine Fever (ASF)**

I would like to move into the national **communication strategy** on ASF, based on your knowledge and experience.

17. Could you please tell me if there is a **department in charge** of the communication as part of the strategy of ASF? *If **Yes**,* Could you please tell me the name of that department?

18. *Which are the **main** target groups of the communication strategy of ASF?

*Potential target groups:*

| | | |
|---|---|---|
| *Smallholder pig keepers* | | *Hunters* |
| *Commercial farmers* | | *Forest workers* |
| *Temporal farm workers* | | *Veterinarians* |
| *Pig-breeders* | | *Transport authorities and check point staff* |
| *Livestock transporters* | | *General public (esp. travellers)* |
| *Slaughterhouse workers* | | |

19. *Which are the **main** communication channels used for those target groups?

20. *What would you describe as the **shortcomings** of the current communication strategy?

    *Why do you say that…?*

    *Shortcoming: deficiency, weak points*

21. Based on what you have told me so far, I would like to know what you think of the use of **social media** when it comes to a national communication strategy on an animal health emergency such as ASF compared to traditional channels (newspapers, magazines, radio, TV)?

    *Why do you say that?*

22. Do you know if social media is **regularly used** to share news/ information related to the current emergencies of ASF in Estonia?

23. *If yes, which social media is being used? Would you say that this communication strategy through social media is effective? Could you rate your perception of the rate of success?*

    *Levels for rating: poor, fair, good, excellent*
    *Please, justify your rating…*

S5 Table: Number of farmers interviewed by county (Estonian region).

| County | Number | Frequency (%) |
|---|---|---|
| Lääne-Viru | 5 | 22.7 |
| Tartu | 3 | 13.6 |
| Jõgeva | 4 | 18.2 |
| Põlva | 1 | 4.5 |
| Harju | 2 | 9.1 |
| Järva | 1 | 4.5 |
| Saare | 5 | 22.7 |
| Ida-Viru | 1 | 4.5 |
| Total | 22 | 100 |

S6 Table: Biosecurity aspects that Estonian farmers mentioned to have invested resources due to ASF.

| Question | Category | Number of responses | Percentage (%) |
|---|---|---|---|
| In what aspects of biosecurity did you invest resources in your farm due to ASF? | Cleaning and disinfection | 18 | 81.8 |
| | Constructional changes | 13 | 59.1 |
| | Equipment supply | 8 | 36.4 |
| | Feed, water | 9 | 40.9 |
| | Fencing | 20 | 90.9 |
| | Management of dead animals | 17 | 77.3 |
| | Other | 1 | 4.5 |
| | Removal of manure | 7 | 31.8 |
| | Sourcing of animals | 6 | 27.3 |
| | Training staff on disease management | 20 | 90.9 |
| | Transport of animals | 17 | 77.3 |

S7 Table: Public institutions and private organizations mentioned by the Estonian veterinary authorities as part of the risk management strategy.

| **Public Institutions** | **Nominations** |
|---|---|
| Veterinary and Food Board | 9 |
| Veterinary and Food laboratory | 7 |
| Environmental Board | 7 |
| Ministry of Rural Affairs | 5 |
| University of Life Sciences | 5 |
| Estonian environmental agency | 3 |
| Ministry of environment | 2 |
| Rendering plant | 2 |
| Tax and custom board | 1 |
| Environmental inspectorate | 1 |
| Police | 1 |
| **Private organizations** | **Nominations** |
| Estonian hunters' association | 9 |
| Pig farmers' association | 4 |
| Private veterinarians | 2 |
| Pig breeder society | 1 |

S8 Table. Target groups, information material and communication channels mentioned by the Estonian veterinary authorities

| Target group | Communication channels | | | |
| --- | --- | --- | --- | --- |
| | **Electronic channels** | **Written methods** | **Face-to-face communication** | **Broadcast media communication** |
| *Smallholder pig keepers* | Website of the ministry of rural affairs, emails, posters, leaflets | Posters, leaflets, newspaper articles | Meetings | |
| *Commercial farmers* | Website of the ministry of rural affairs, emails | Posters, leaflets, newspaper articles | Meetings | |
| *Temporal farm workers* | Website of the ministry of rural affairs | | | |
| *Pig-breeders* | Website of the ministry of rural affairs, posters, leaflets | Posters, leaflets, newspaper articles | | |
| *Livestock transporters* | Website of the ministry of rural affaires | | | |
| *Slaughterhouse workers* | Website of the ministry of rural affairs | | Oral presentations of official veterinarians to workers | |
| *Hunters* | Website of the ministry of rural affairs, posters, leaflets, emails with information updated for the district | Magazine from the hunter’s association, posters, leaflets, newspaper articles | Meetings with the veterinary and food board | |
| *Forest workers* | Website of the ministry of rural affairs | | | |
| *Veterinarians* | Website of the ministry of rural affairs | | | |
| *Transport authorities and check point staff* | Website of the ministry of rural affairs, posters | Posters | | |
| *General public* | Website of the ministry of rural affairs, posters | Posters, newspapers articles | | News on the TV/radio, Facebook page of the veterinary and food board |
| *Military forces/government rescue service* | Website of the ministry of rural affairs, NETO guidelines about biosecurity rules in the environment | | Military events | Public news about biosecurity to soldiers |

| | | | |
|---|---|---|---|
| *Feed producers* | Website of the ministry of rural affairs, emails | | |
| *Official vets in meat plants* | | | Training courses |

S9 GDELT Query for ASF news

The data were collected using from the GDELT database using Google Bi Query ( https://cloud.google.com/bigquery) through the following query.

```
SELECT DocumentIdentifier, DATE, V2THEMES, V2Organizations, V2Persons, V2Locations,
TranslationInfo
FROM `gdelt-bq.gdeltv2.gkg_partitioned`
WHERE _PARTITIONTIME >= "2015-01-01 00:00:00"
AND ((DocumentIdentifier LIKE '%swine%fever%') or (DocumentIdentifier LIKE
'%influenza%suina%')
 or (V2THEMES LIKE '%TAX_DISEASE_AFRICAN_SWINE_FEVER%'))
```
